# Inhibition of VCP preserves retinal structure and function in autosomal dominant retinal degeneration

**DOI:** 10.1101/2020.11.17.384669

**Authors:** Blanca Arango-Gonzalez, Merve Sen, Rosellina Guarascio, Kalliopi Ziaka, Eva M. del Amo, Kwan Hau, Hannah Poultney, Rowan Asfahani, Arto Urtti, Tsui-Fen Chou, Sylvia Bolz, Raymond J. Deshaies, Wadood Haq, Michael E. Cheetham, Marius Ueffing

**Author notes:** Corresponding author: Marius Ueffing, address: Centre for Ophthalmology, Institute for Ophthalmic Research, University of Tuebingen, Elfriede-Aulhorn-Str. 7, D-72076 Tuebingen, Germany. Co-first authors (Blanca Arango-Gonzalez, Merve Sen).

## Abstract

Due to continuously high production rates of rhodopsin (RHO) and high metabolic activity, photoreceptor neurons are especially vulnerable to defects in proteostasis. A proline to histidine substitution at position 23 (P23H) leads to production of structurally misfolded RHO, causing the most common form of autosomal dominant Retinitis Pigmentosa (adRP) in North America. The AAA-ATPase valosin-containing protein (VCP) extracts misfolded proteins from the ER membrane for cytosolic degradation. Here, we provide the first evidence that inhibition of VCP activity rescues degenerating P23H rod cells and improves their functional properties in P23H transgenic rat and P23H knock-in mouse retinae, both *in vitro* and *in vivo*. This improvement correlates with the restoration of the physiological RHO localization to rod outer segments (OS) and properly-assembled OS disks. As a single intravitreal injection suffices to deliver a long-lasting benefit *in vivo*, we suggest VCP inhibition as a potential therapeutic strategy for adRP patients carrying mutations in the *RHO* gene.

## INTRODUCTION

Retinitis pigmentosa (RP) is a group of inherited vision disorders causing progressive and irreversible degeneration of retinal photoreceptor cells. More than 3,000 mutations in over 70 different genes are known to cause non-syndromic RP, and the autosomal dominant form (adRP) accounts for approximately 15–25% of cases (1). In turn, over 150 different mutations in the rhodopsin *(RHO)* gene have been identified, and collectively, they are the most common cause of adRP (2, 3). This gene encodes for rhodopsin (RHO) that is expressed by rod photoreceptors and makes up > 80 % of all proteins in the disk membranes of the rod photoreceptor outer segments (OS) (4–6). The variant *RHO_P23H_* is the most common cause of adRP in the United States and is the most extensively studied RHO mutation leading to misfolding (7). Structurally misfolded RHO^P23H^ is retained within the ER of the inner segment (IS) of rod photoreceptor cells, where RHO is synthesized and degraded by ER-associated degradation (ERAD) (8).

ERAD removes misfolded proteins that fail to achieve their native state (9) and thereby contributes to proteostasis. In the retina, balanced proteostasis is critical for cell survival, and its imbalances for prolonged periods can result in cell death (2). The *RHO_P23H_* mutation can lead to an accumulation of misfolded mutant protein in the ER of photoreceptors and to activation of the unfolded protein response (10), potentially with insufficient or imbalanced unfolded protein response output (11). In the long run, the presence of RHO^P23H^ results in a metabolic imbalance, mitochondrial failure, and ultimately selective photoreceptor cell degeneration (12).

Transgenic, knock-out, and knock-in (KI) animal models allow scientific investigation of retinal degeneration in RP. P23H transgenic rats carry multiple copies of the variant *RHO_P23H_* allele and are widely used as an animal model for adRP (13). The P23H KI mouse model closely reflects the genetic load found in adRP patients. In both animal models, the retinae display progressive rod photoreceptor degeneration followed by secondary cone degeneration, a phenotype consistent with the clinical findings in patients carrying the *RHO_P23H_* mutation (14–17).

The ATPase valosin-containing protein (VCP) is one of the most abundant cytosolic proteins. It is mainly localized in the cytosol, with a significant fraction associated with organelle membranes, such as the ER and the Golgi (18). In photoreceptors, VCP, also known as p97, specifically acts as a quality control checkpoint for membrane proteins (12). VCP is an AAA+ATPase with two ATPase domains (D1 and D2) and is equipped with an N-terminal domain that recruits cofactor/substrate specificity factors. VCP forms a functional hexamer that acts as an ATP-dependent molecular machine (19, 20). It extracts proteins from macromolecular complexes or membranes resulting in downstream activity, unfolds ubiquitylated proteins for proteasomal degradation, and also functions as an interaction hub, with more than 30 cofactors able to modulate VCP-mediated processes (21–24). The ATP hydrolyzing activity of VCP is indispensable for its function, and ATP consumption is required to extract incorrectly folded proteins from the membranes of the ER or mitochondria (25).

*VCP* mutations have been associated with several diseases. Point mutations have been linked to multisystem proteinopathy (MSP), also known as IBMPFD (Inclusion Body Myopathy associated with Paget’s disease of the bone and Frontotemporal Dementia) and amyotrophic lateral sclerosis (ALS) (21, 26–30). In these conditions, the dysfunction of VCP is considered to result in the deregulation of the proteostasis network (26, 31). Conversely, excessive ATPase activity of VCP is also pathogenic (32), resulting in mitochondrial fragmentation, cell death in neurons (33), and photoreceptor cell degeneration (34).

In an earlier study, we showed that VCP binds to both normal RHO as well as RHO^P23H^ before its release from the ER when expressed in HEK-293 cells (35). Moreover, genetic inactivation of VCP in *Drosophila* carrying the mutant *RHO^P37H^* allele (*Rh1^P37H^*) suppressed retinal pathology (36). Here, we examined whether modulation of the ER-associated degradation machinery by VCP inhibition could have a therapeutic impact on mammalian photoreceptors expressing RHO^P23H^ in two rodent models for adRP.

## RESULTS

VCP unfolds and extracts misfolded proteins from the ER membrane in an ATP-dependent manner. To investigate the effect of inhibition of this highly energy-consuming step in mammalian photoreceptors, we treated retinae from P23H transgenic rats and P23H KI mice with VCP inhibitors. We selected three chemically unrelated small-molecule VCP inhibitors, ML240, Eeyarestatin I (EerI), and NMS-873. ML240 acts as a selective and reversible ATP-competitive inhibitor of VCP, targeting its D2-domain (20, 37, 38). EerI acts as an irreversible inhibitor binding the D1 domain of VCP. EerI influences deubiquitinating processes mediated by VCP-associated enzymes, but does not inhibit the ATPase activity of purified VCP (38–40). NMS-873 is an allosteric ATP- non-competitive VCP inhibitor that binds at the D1–D2 interdomain linker and stabilizes the D2-ADP-bound form, highly potent and with high selectivity. (19, 31).

### VCP inhibition rescues rod photoreceptors in P23H transgenic rat explanted retinae

We first tested two VCP inhibitors (ML240 and EerI) in a serum-free organotypic culture system that preserves the retinal pigment epithelium (RPE) attached to the distal part of the OS (41). Organotypic retinal cultures allowed us to test both VCP inhibitors at different doses *in vitro* under standardized conditions. In previous studies, we have shown that photoreceptor degeneration in P23H-1 rats starts around postnatal day (PN) 9 and peaks at PN15 (14, 42), see Fig. S1. Accordingly, P23H retinae were isolated from PN9 and cultivated for six days (Fig. S1). Within this time, photoreceptor cell death and regression of the outer nuclear layer (ONL) resembled the process of retinal degeneration *in vivo*. Organotypic cultures were treated every second day with ML240 or EerI at different doses in order to select the most effective concentration, based on cell survival in the ONL (data not shown). Since both compounds (ML240 and EerI) require DMSO as vehicle, we selected the lowest amount of DMSO that allowed complete solubilization of each compound; thus, corresponding vehicle-control groups were treated with 1 % or 0.5 % DMSO, respectively.

To identify photoreceptors undergoing cell death, we used the TUNEL assay (Fig. 1A), which allowed us to determine the percentage of TUNEL-positive dying cells compared to the total number of ONL cell nuclei. Both ML240 and EerI significantly reduced the percentage of photoreceptor cell death in P23H cultured retinae compared to the respective controls (Vehicle: 6.27 % ± 1.91; ML240: 2.23 % ± 0.36 p<0.0001; Vehicle: 5.08 % ± 0.41; EerI: 1.70 % ± 0.39 p=0.0004 (Fig. 1B)). This reduction in cell death was reflected in an increased number of remaining cell rows in the ONL. Thus, we found that the ONL of ML240 and EerI treated P23H retinae contained significantly more rows of nuclei compared to the respective controls (Vehicle: 7.75 rows ± 0.76; ML240: 9.49 rows ± 0.93 p=0.0026; Vehicle: 8.08 rows ± 0.54; EerI: 9.48 rows ± 0.21 p=0.0344 (Fig. 1C)).

**Figure 1.**
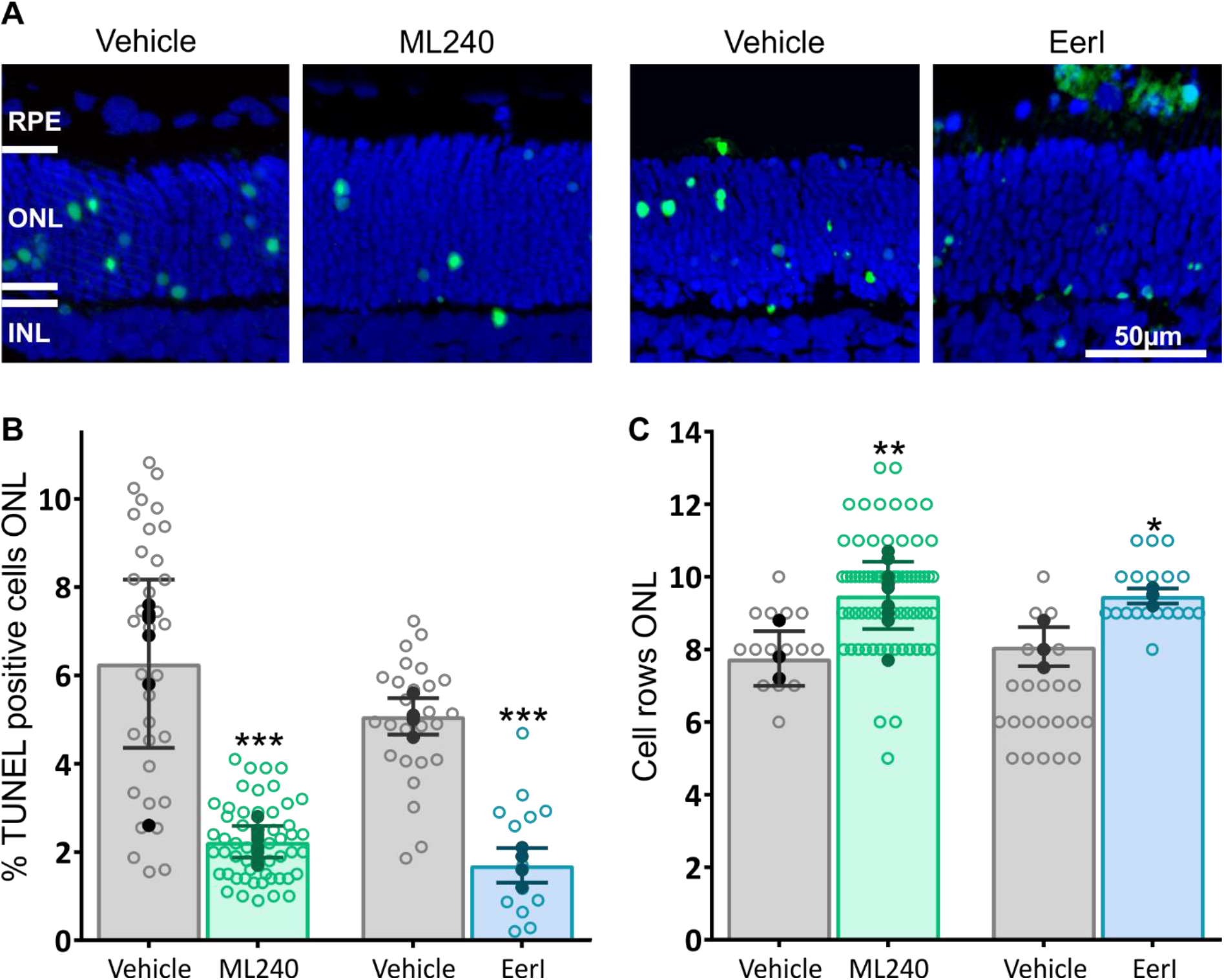
Neuroprotective effect of VCP inhibition in organotypic cultures of P23H retinae. Retinae from P23H transgenic rats were explanted at postnatal day 9 (PN9), kept for 6 days *in vitro* (DIV6), and treated every second day with either 20 μM ML240 or 10 μM Eeyarestatin (VCP inhibitors). ML240 and EerI were dissolved in the smallest practical volume of DMSO, and corresponding controls were treated using the same vehicle volume. **(A)** Explants were stained with TUNEL assay to differentiate photoreceptors undergoing cell death (green) using nuclei counterstaining with DAPI (blue). **(B)** Bar chart shows the percentage of TUNEL-positive cells in the ONL. A significant decrease in the percentage of dying cells was observed after treatment with either ML240 or EerI compared to the respective vehicle controls. Comparison of the number of remaining cell rows in the ONL. Retinae treated with VCP inhibitors showed significant preservation of the number of photoreceptor cell rows. Values were quantified by scoring several images (open circles) from at least four retinae (closed circles, n=4) per treatment. Data plotted as mean ± SD. One-way ANOVA with Bonferroni multiple comparison test; ***p<0.001; **p<0.01, *p<0.05. RPE: retinal pigment epithelium; ONL: outer nuclear layer; INL: inner nuclear layer. Scale bar: 50μm.

### Single intravitreal injection of VCP inhibitors to P23H transgenic rats protects degenerating rod photoreceptors *in vivo*

Next, we evaluated the effect of VCP inhibition in P23H transgenic rats *in vivo*. To avoid potential unwanted systemic effects of VCP inhibition and to be as close as possible to clinically approved drug delivery procedures into the eye, we administered both compounds (ML240 and EerI) by intravitreal injection. We selected PN10 as the injection day and PN15 as our first evaluation time-point, where the peak of degeneration in P23H transgenic rats has been observed (Fig. S1). This group was designated the short-term (ST) group. Similar to our observations *in vitro*, five days after the injection of VCP inhibitors in the ST group, the TUNEL assay indicated a significant decrease in the percentage of dying photoreceptor cells with either ML240 or EerI (Vehicle: 2.76 % ± 1.30; ML240: 1.20 % ± 0.66 p=0.001; Vehicle: 2.68 % ± 0.64; EerI: 0.47 % ± 0.27 p<0.0001 (Fig. 2A and B)). Concomitantly, we observed a statistically significant increase in the number of remaining cell rows in the ONL in the EerI treated retinae during this short period (Vehicle: 9.56 rows ± 0.59; ML240: 9.75 rows ± 0.54 p>0.99; Vehicle: 10.28 rows ± 0.51; EerI: 11.57 rows ± 0.72 p=0.0003 (Fig. 2C).

**Figure 2.**
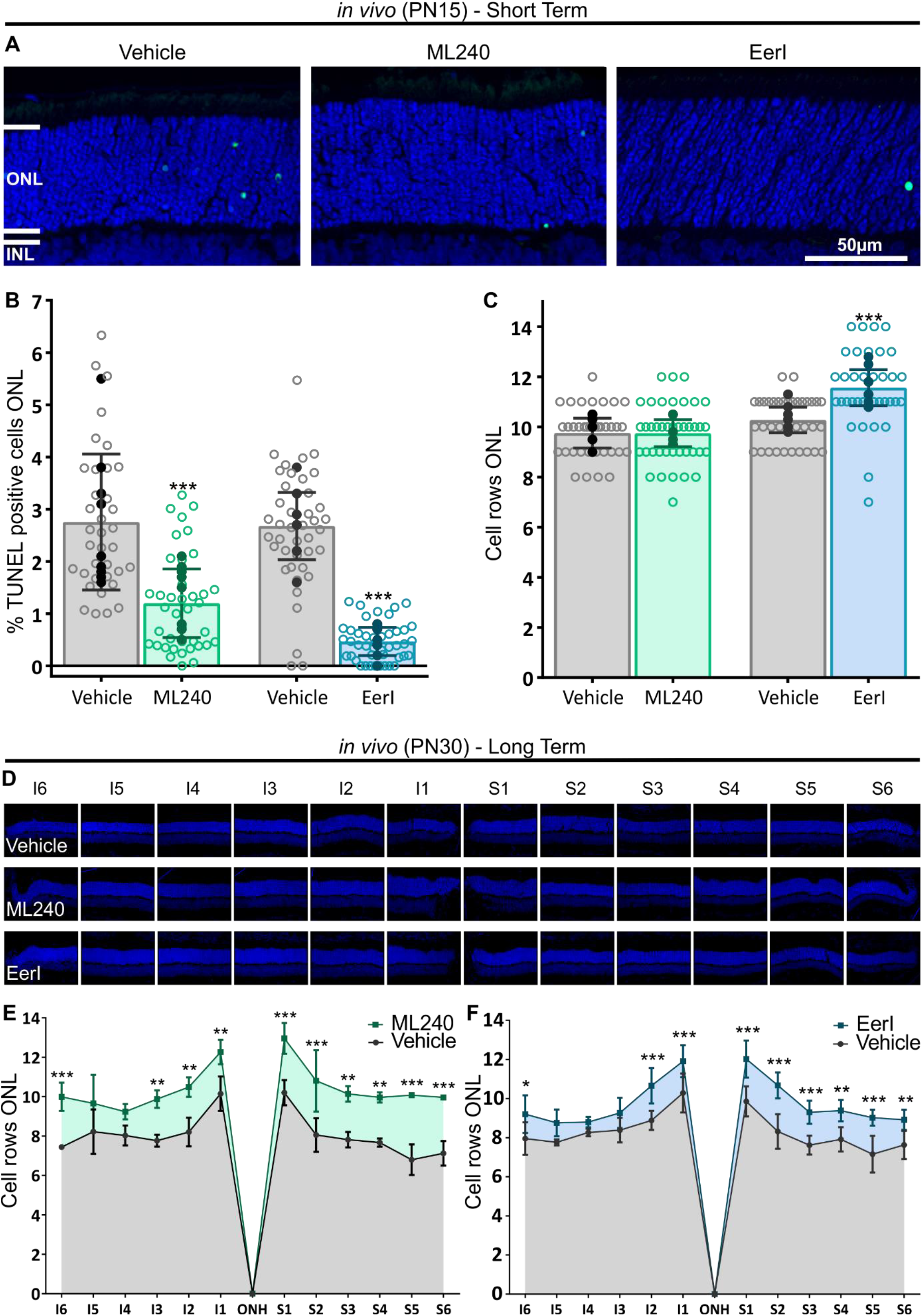
Single intravitreal injection of VCP inhibitors to P23H transgenic rats protects degenerating rod photoreceptors *in vivo*. Intravitreal injections in P23H rats were performed at postnatal day 10 (PN10). For the analysis, the animals were divided into short-term (retinae fixed at PN15, A-C) and long-term (retinae fixed at PN30 D-F). **(A)** Representative images of the stained retinae fixed at PN15 using the TUNEL assay to differentiate photoreceptors undergoing cell death (green) and nuclei counterstaining using DAPI (blue). **(B)** Bar chart shows the percentage of TUNEL-positive cells in the PN15 fixed retinae. In the short term treated group, both ML240 and EerI significantly reduced the number of dying cells compared to the respective vehicle control (contralateral retina) in the P23H mutant. **(C)** Quantification of the ONL cell rows for the PN15 fixed retinae. Only in the EerI treated group a significant increase in the number of remaining cell rows in the ONL 5 days after treatment was observed. **(D)** Representative images of the radial parasagittal sections showing inferior (I) and superior (S) hemispheres of the retinae stained with DAPI (blue) of vehicle, ML240, and EerI long-term treated groups. **(E and F)** Retinal spidergrams of the inferior and superior hemispheres along the vertical meridian revealed that treated retinae (E: ML240 in green, F: EerI in blue) contain more nuclei rows in the ONL than the contralateral vehicle-treated retinae. The neuroprotective effect is stronger in the superior retina. Values were quantified by scoring several images (open circles) from at least four retinae (closed circles, n=4) per treatment. Plotted data as mean ± SD. One-way ANOVA with Bonferroni multiple comparison test; (B-C) or Two-way ANOVA with Bonferroni multiple comparison test; (E-F). ***p<0.001; **p<0.01; *p<0.05. ONL: outer nuclear layer; INL: inner nuclear layer; ONH: optic nerve head. Scale bar: 50μm.

To estimate the duration of the protective effect, we examined a long-term (LT) group that was injected at PN10 and euthanized at PN30 (Fig. S1). We selected this age because rat retinae and their photoreceptors reach a mature development level around the first postnatal month. We evaluated the number of remaining photoreceptor cell rows in the ONL at six locations across both the retina’s inferior and superior hemispheres. This is because the P23H transgenic rat retinae at P30 show a higher degree of photoreceptor degeneration in the superior hemisphere than in the inferior hemisphere (15) (Fig. 2D-F). Even 20 days after a single injection, we could detect a significant neuroprotective effect of both ML240 and EerI (Fig. 2E and F, and Table S1). This sustained effect was stronger and more pronounced in the superior retina.

To estimate the intravitreal clearance (CL_ivt_) of the two VCP inhibitors, we performed model calculations based on the Quantitative Structure-Property Relationships (QSPR) model, that uses comprehensive rabbit data from intravitreal pharmacokinetic experiments (43). Assuming similar permeabilities in rat and rabbit RPE, the CL_ivt_ (CL = P x S, where P is the permeability and S is the RPE’s surface area) in the rat would be 0.04 ml/h. By using the equation: t_1/2_ = ln2 Vd/CL_ivt_ (t_1/2_ is the half-life, Vd: is the volume of distribution estimated to be similar to the anatomical volume of the vitreous (44), of young rats, 25 μl (45)), we calculated the expected half-life in the vitreous to be approximately 25 minutes for both compounds. The simulated concentration profiles of the *in vivo* treatment with ML240 and EerI in the rat vitreous are presented in Fig. 3, showing more than 98 % of the dose is expected to be eliminated within the first 3 hours post-injection. Nevertheless, both VCP inhibitors were still protective at PN30, 20 days after injection. This sustained protective effect suggests that the mechanisms promoting photoreceptor cell survival started by VCP inhibition remain active even after most, if not all, of the VCP inhibitor, has already been eliminated from the vitreous.

**Figure 3.**
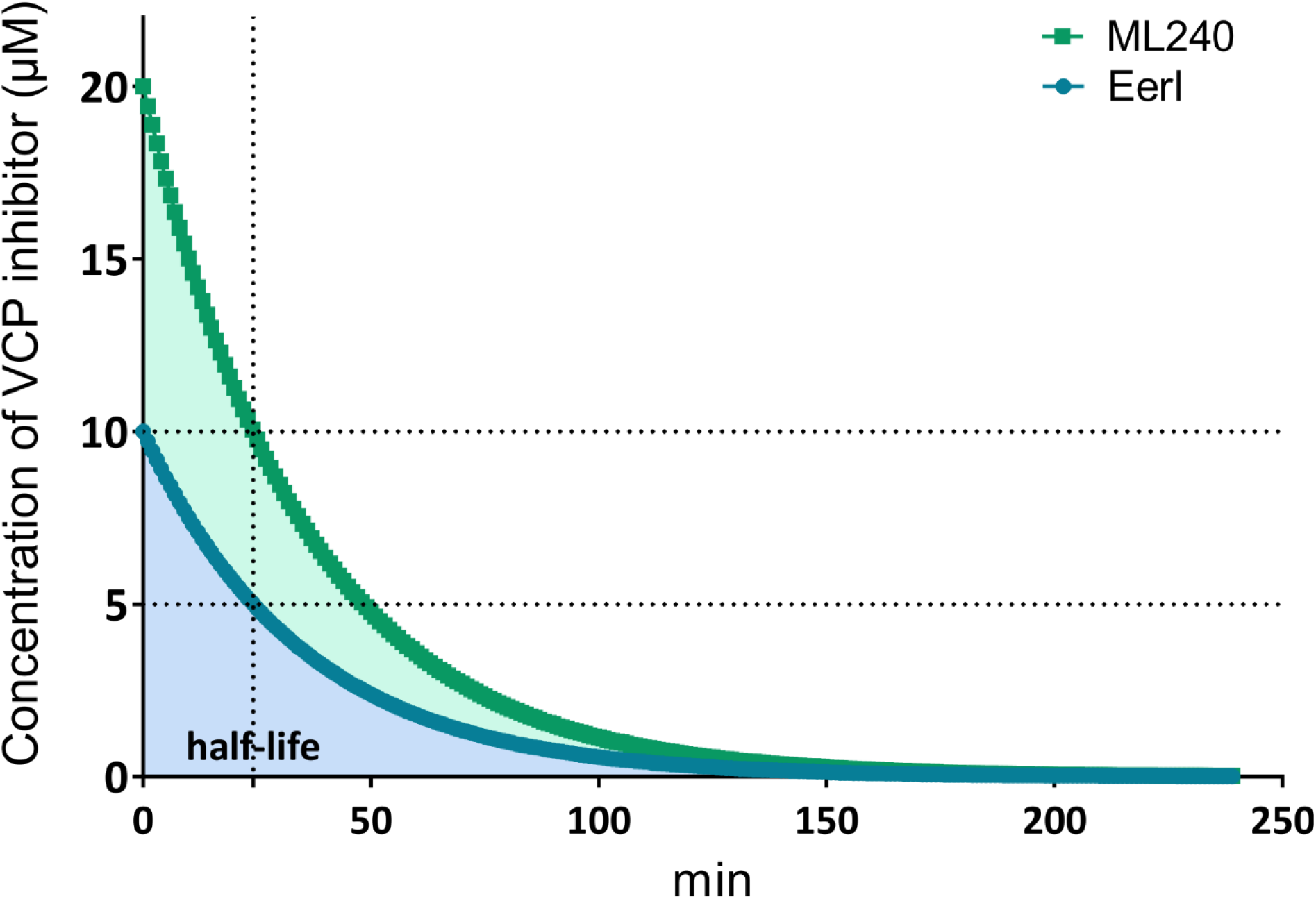
Simulated concentration profiles of ML240 and Eerl. After intravitreal injection of 0.5 nmol and 0.25 nmol dose respectively in the rat vitreous, using the estimated rat pharmacokinetic parameters of intravitreal clearance 0.04 ml/h and volume of distribution 25 μL for both analyzed groups, the expected half-life was calculated to be approximately 25 minutes; thus, more than 98 % of both drugs is expected to be eliminated in the first 3 hours after injection. Since ML240 and EerI were still protective 20 days after the single injection, mechanisms promoting photoreceptor cell survival started by VCP inhibition presumably remain active even after most of the drug has already been eliminated from the vitreous.

### VCP inhibition restores RHO localization and increases the OS length in P23H transgenic rat retinae *in vitro* and *in vivo*

Some of the features of P23H photoreceptor degeneration are an increase in RHO staining in the ONL and IS and the perturbation of the OS structure (17, 46, 47). Therefore, we analyzed the effect of VCP inhibition on RHO expression and localization by immunofluorescence staining assessing three different parameters: (1) distribution of RHO immunostaining in the retina, (2) mean peak of fluorescence intensity in the somata (ONL), and (3) length of the OS.

We used WT rat retinae at PN15 and PN30 as controls. Here, and as previously described (48), RHO immunostaining localizes mainly at the OS (Fig. 4A and C, PN15, and P30, respectively). In contrast, RHO was abnormally distributed in P23H rod photoreceptor cells, similar to observations made in other animal models for RP (49–51). Here, RHO accumulated in both the ONL and IS of the photoreceptor cells with a reduced distribution to the OS (Fig. 4A and C, PN15, and PN30, respectively).

**Figure 4.**
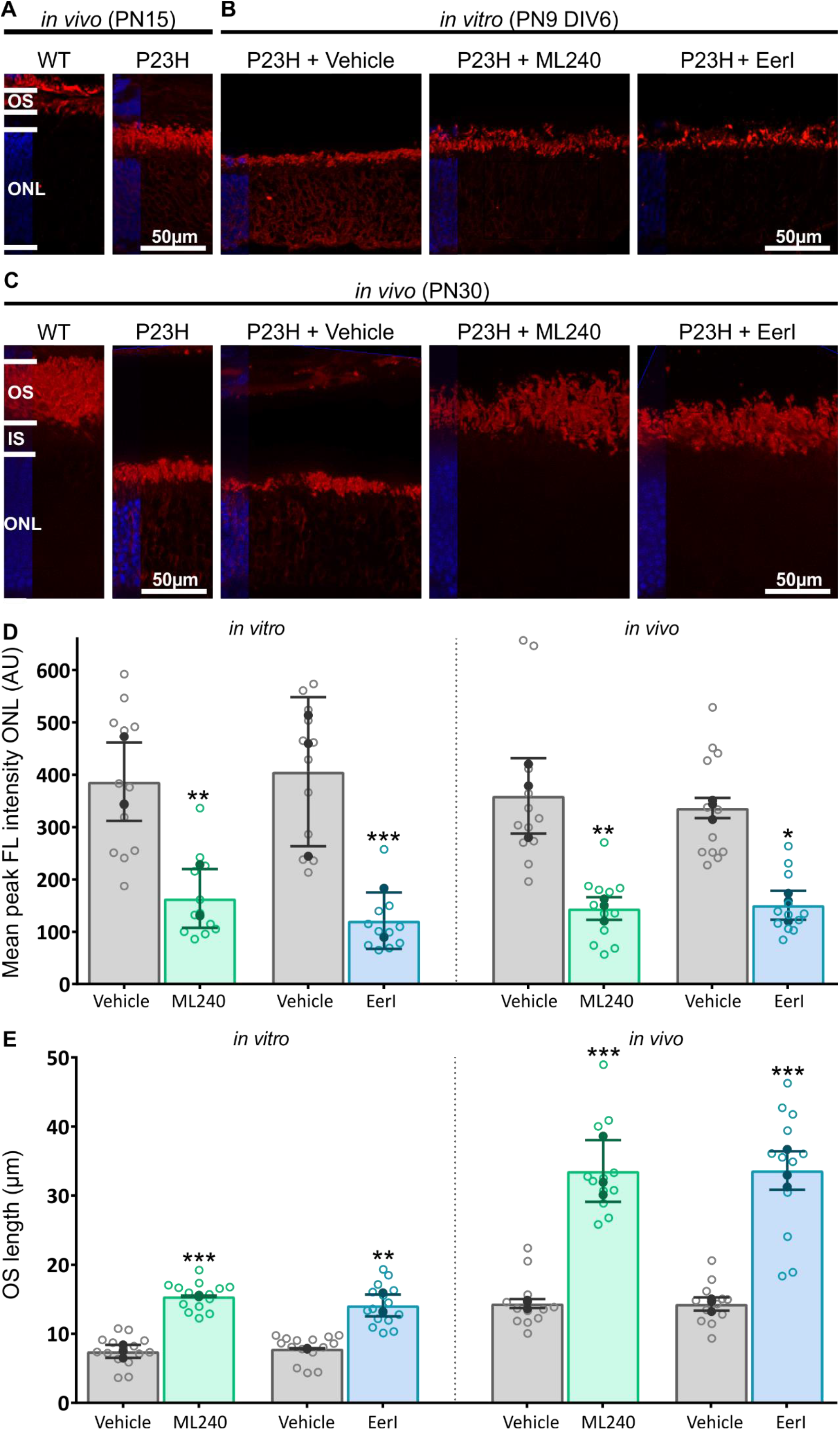
VCP inhibition restores RHO localization and increases OS length in P23H transgenic retinae *in vitro* and *in vivo*. Immunofluorescence labeling in cryosections designates the location of RHO (red staining) in **(A)** Untreated WT and P23H control retinae at PN15; **(B)** organotypic cultures of P23H retinae, explanted at postnatal day 9 and cultivated for 6 days (PN9 DIV6), treated with vehicle, ML240, or EerI; and **(C)** PN30 *in vivo* retinae from untreated WT and P23H and treated P23H rats (after a single injection at PN10 of vehicle, ML240, or EerI). Untreated WT retinae at PN15 and PN30 as a point of reference, as staining in WT, was exclusively observed in the OS. In contrast, in P23H untreated retinae, RHO staining was observed in both the ONL and the OS, indicating an accumulation in the mutants’ cell somata. In the vehicle control groups, *in vitro,* as well as *in vivo*, an abnormal RHO distribution was observed, with strong fluorescence staining in the ONL and very short rod OS. In retinae treated with VCP inhibitors - *in vitro* and *in vivo* we found a reduction of the fluorescence in the ONL and better preserved and longer OS. VCP inhibitors were able to restore the distribution of RHO predominantly to the OS and increased OS length. **(D)** Quantification of RHO immunofluorescence intensity in the ONL *in vitro* and *in vivo*. A central region of each image was selected, and the mean maximum intensity was assessed using the Zen 2.3 software. *In vitro*, as well as *in vivo*, VCP inhibition reduced the FL intensity in the ONL. **(E)** The mean length of the OS was significantly higher in both *in vitro* and *in vivo* treated retinae when compared to the vehicle controls. Values were quantified by scoring several images (open circles) from three retinae (closed circles, n=3) per treatment. Data plotted as mean ± SD. One-way ANOVA with Bonferroni multiple comparison test; ***p<0.001; **p<0.01; *p<0.05. OS: outer segments; IS: inner segments; ONL: outer nuclear layer; AU: fluorescence arbitrary-units. Scale bar: 50μm.

To test whether VCP inhibition can correct the decrease of RHO in OS, we evaluated RHO immunostaining of P23H retinae treated with either ML240 or EerI in retinal organ cultures (*in vitro*) or the intact eye after intravitreal injections (*in vivo*). In both sets of experiments (*in vitro* and *in vivo*), the RHO staining in vehicle-control P23H retinae was mislocalized, similar to the untreated retinae (Fig. 4B and C).

In contrast, VCP inhibition in P23H retinae (*in vitro* or *in vivo*) almost completely restored RHO’s distribution to that of the normal WT phenotype, with staining predominantly in the OS. (Fig. 4B, and C). This WT-like distribution suggests that either RHO was successfully trafficked from the ER to OS disks or there was enhanced removal of the mutant protein and reduction in any dominant-negative effects, unlike in the untreated P23H retinae. To measure RHO distribution, we quantified the mean peak of fluorescence intensity in the ONL (Fig. 4D). To do so, we selected the ONL area in images taken under identical conditions and calculated the fluorescence intensity and standard deviation (SD). The observed significant decrease of the mean peak of RHO fluorescence intensity in the treated retinae for both *in vitro* (Vehicle: 386.70 ± 74.65; ML240: 163.60 ± 56.18 p=0.0048; Vehicle: 405.9 ± 142.4; EerI: 121.10 ± 53.86; p=0.0005; Fig. 4D) and *in vivo* (Vehicle: 359.80 ± 72.09; ML240: 144.50 ± 21.68; p=0.0064; Vehicle: 336.50 ± 19.31; EerI: 150.70 ± 27.60; p=0.0191 (Fig. 4D)), confirmed the reduced retention of RHO in the ER. Also, measurements of OS length supported the hypothesis that there was improved traffic to the OS (Fig. 4E). OS of retinae treated with ML240 or EerI were significantly longer, both *in vitro* (Vehicle: 7.45 μm ± 0.95; ML240: 15.42 μm ± 0.11; p=0.0007; Vehicle: 7.85 μm ± 0.06; EerI: 14.10 μm ± 1.59; p=0.0063; Fig. 4E) and *in vivo* (Vehicle: 14.38 μm ± 0.65; ML240: 33.56 ± 4.47; p<0.0001; Vehicle: 14.31 μm ± 0.96; EerI: 33.64 μm ± 2.79; p<0.0001; Fig. 4E). As RHO composes the vast majority of the total OS integral membrane protein (6), the reduction of RHO staining in the ONL accompanied by an increase of the OS length strongly suggest that RHO is no longer retained in the IS once VCP is inactivated, allowing proper membrane localization and translocation to the OS.

At this point, we found similar protective characteristics between ML240 and EerI. To more efficiently use laboratory animals and following the 3Rs - reducing, refining, and replacing animal use (52), we decided to select only one of the compounds for further experiments. Inhibition with EerI is irreversible and unspecific (inhibition of other cellular components, including ataxin 3 and Sec61), and it has been suggested that EerI needs to be metabolized into an active compound to exert its inhibitory effect (31, 39, 53). In contrast, VCP inhibition by ML240 works by a defined mechanism of action, is reversible, and ML240 has low off-target activity compared to EerI. Within the context of the Ubiquitin/Proteasome System, ML240 is specific for VCP, is far-better characterized in terms of VCP-dependent processes, and selectively inhibits the D2 ATPase domain of VCP (19, 20, 38). For these reasons, we focused on VCP inhibition by ML240 for the ultrastructure analysis and the retinal function evaluation in the P23H rat model of RP.

### VCP inhibition corrects P23H destabilization in rod disk membranes in P23H rats

We used electron microscopy (EM) to analyze how the improvement in the RHO localization and cell survival due to VCP inhibition affected OS’s ultrastructural properties, especially concerning OS disk structure and organization. Again, we included both *in vitro* and *in vivo* paradigms. For *in vitro* assessment, P23H retinae were explanted at PN20 and treated with ML240 for two days to evaluate the acute effect of VCP inhibition. For *in vivo* assessment, we intravitreally injected ML240 at PN10 and analyzed the eyes at PN21 (Fig. S1). We selected this time-point as the photoreceptor structure has fully matured after PN20 (54). Reynold’s lead citrate-stained semi-thin sections of the treated explants showed longer and morphologically more homogeneously structured OS than vehicle controls (Fig. 5A and B).

**Figure 5.**
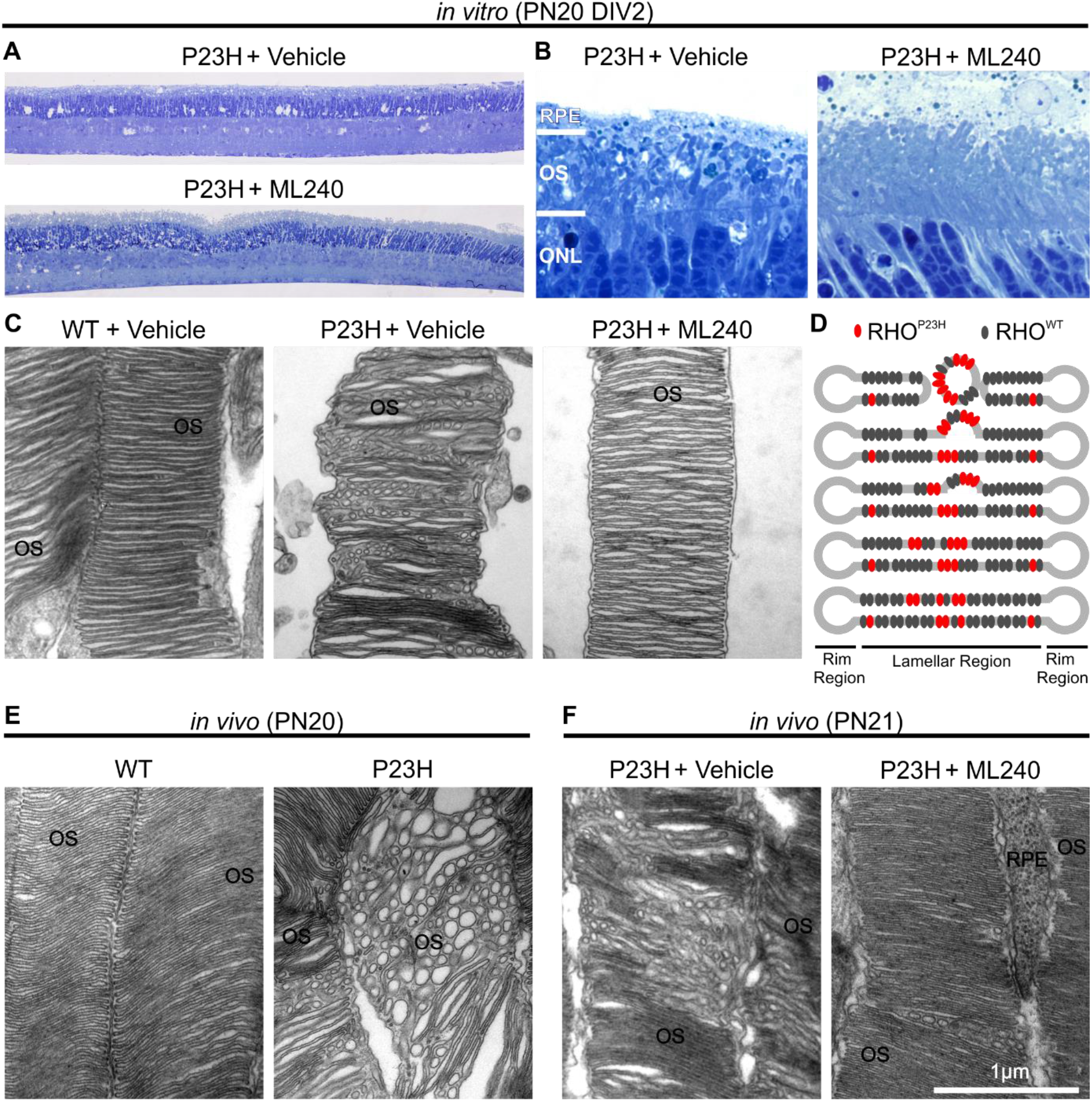
VCP inhibition corrects P23H destabilization in rod disk membranes. Ultrastructure analysis of longitudinal sections of rod photoreceptor cells using transmission electron microscopy (EM) in WT and P23H rat retinae *in vitro* and *in vivo*. **(A and B)** Reynold’s lead citrate-stained semithin sections of vehicle control and ML240 treated retinae (postnatal day 20 and cultivated for 2 days (P20DIV2)) show increased preservation of the outer retina and especially of the OS after VCP inhibition. **(C)** EM of OS from organotypic cultures (PN20 DIV2) of WT and P23H retinae treated with vehicle or ML240. OS of vehicle-treated P23H cultures exhibited numerous vesicotubular structures similar to those described in Haeri and Knox, 2012 (55). **(D)** Schematic model for defective disk formation in OS in the P23H mutant (adapted from (55)). In this model and depending on a critical concentration, P23H mutant RHO (red dots) begins to self-associate and form aggregates in the membrane, excluding WT protein (grey dots). Thus, mutant protein concentrated in a localized area causes defects in the membrane structure, leading to vesiculation followed by disk breakdown. **(E)** EM of OS from WT and P23H untreated retinae *in vivo*. Only mutant OS display vesicotubular structures. **(F)** EM of OS from P23H of vehicle or ML240-treated retinae *in vivo* via intravitreal injection. Intravitreal injections in P23H rats were performed at postnatal day 10 (PN10) and analysed at PN21 (medium-term, as described in Supplementary figure 1). RPE: retinal pigment epithelium; OS: rod outer segments; ONL: outer nuclear layer. Scale bar: 1μm.

EM examination in WT rat retinae displayed OS discs consisting of a double lamellar membrane connected by a rim region (5) (Fig. 5C and E). *In vitro* or *in vivo* control P23H retinae (untreated or treated with vehicle) revealed abnormal OS disk membranes displaying multiple vesicotubular structures similar to those described in *rho^P23H^*-EGFP transgenic *Xenopus* and *Rho^P23H^* transgenic mice (55), accompanied by broken disk bilayers (Fig. 5C – F). Retinae treated with ML240 showed strongly reduced vesiculation and less ruptured discs in organotypic culture as well as in intravitreally treated eyes (Fig. 5C and F). We calculated the percentage of OS displaying a correct-structured lamellation in ML240 treated retinae compared to vehicle-treated controls. We observed an almost 3-fold increase *in vitro* (Vehicle: 21.88 % ± 1.78; ML240: 59.44 % ± 7.92; p=0.0010) and a significant increase also *in vivo* (Vehicle: 52.42 % ± 11.68; ML240: 84.66 % ± 7.91; p=0.0025; Fig. 6) after treatment.

**Figure 6.**
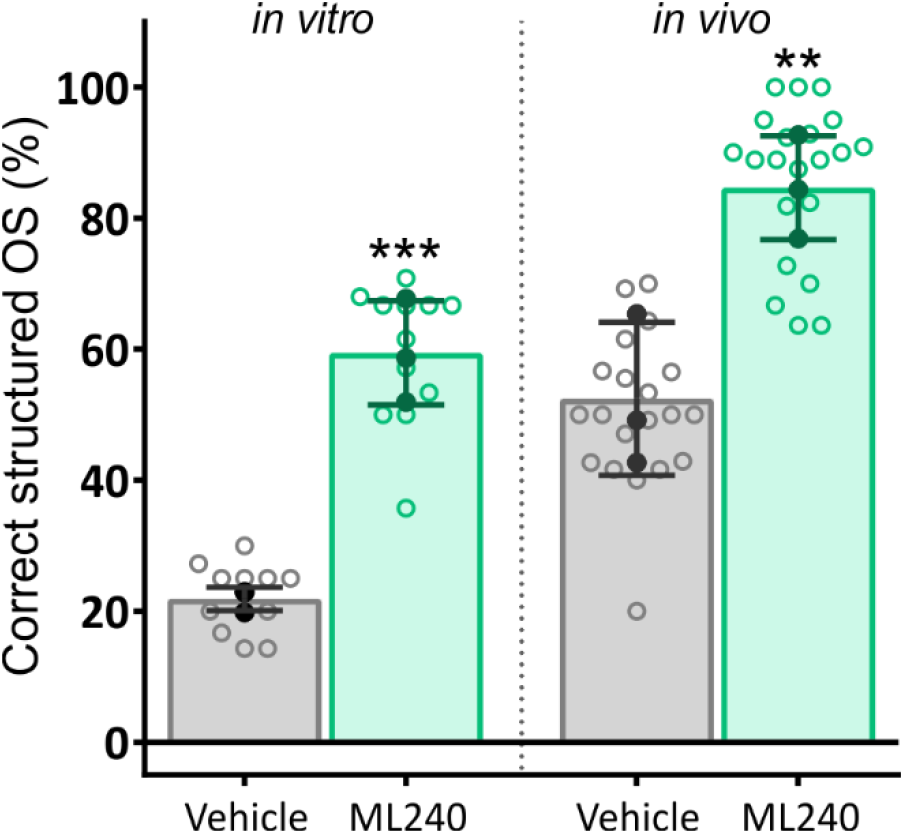
VCP inhibition structurally corrects OS disks in P23H retinae. Graph showing the percentage of correctly structured OS of rod photoreceptor cells using transmission electron microscopy (EM) in P23H rat retinae *in vitro* (PN20 DIV2) and *in vivo* (injected at PN10 and analyzed at PN21, medium-term, as described in Supplementary figure 1). After treatment, in both cultured and *in vivo* injected groups, the percentage of correctly structured OS in EM sections indicated that ML240 treatment significantly preserved the OS structure. Values were quantified by scoring several images (open circles) from at least four retinae (closed circles, n=3) per treatment. Data plotted as mean ± SD; One-way ANOVA with Bonferroni multiple comparison test; ***p<0.001; **p<0.01.

### ML240 treatment results in an improvement of retinal function in P23H rats

To investigate whether the observed preservation of photoreceptor cell viability and structure and improved RHO distribution in P23H retinae after VCP inhibition resulted in improved retinal function, we assessed the retina’s light responsiveness *in vitro* and *in vivo*.

Using a Multi-electrode array (MEA) system, we evaluated the response to light stimulation of PN20 retinae explanted and kept *in vitro* for two days. These organotypic cultures were treated with ML240 and compared to vehicle-treated controls. We positioned the explants on the MEA, excited them using blue light flashes, and recorded the resulting retinal activity (Fig. 7A). Light responsiveness of ML240 treated retinae was significantly increased when compared to vehicle-treated controls. In ML240 treated explants, the percentage of electrodes detecting light-induced activity was approximately six times higher for the ganglion cell activity (Spikes: Vehicle: 9.69 ± 5.33; ML240: 59.75 ± 22.33; p<0.001 (Fig. 7B)) with an activated ganglion cell type distribution of 51.48 % ON, 11.33 % OFF and 37.20 % ON-OFF (Fig. 7C). Along with this, the micro electroretinogram (mERG) indicated a response of more than 80 % of the electrodes in treated retinae and an almost absent response in the control group (mERG: Vehicle: 0.97 ± 1.65; ML240: 84.02 ± 11.32; p<0.0001; Fig. 7B). We also evaluated the retinal explants’ light responsiveness to 20 repetitive light flashes and found that treated explants showed a more than three-fold increased average light response per recording electrode (Vehicle: 2.51 ± 1.96; ML240: 9.28 ± 5.65; p<0.0001; Fig. 7D).

**Figure 7.**
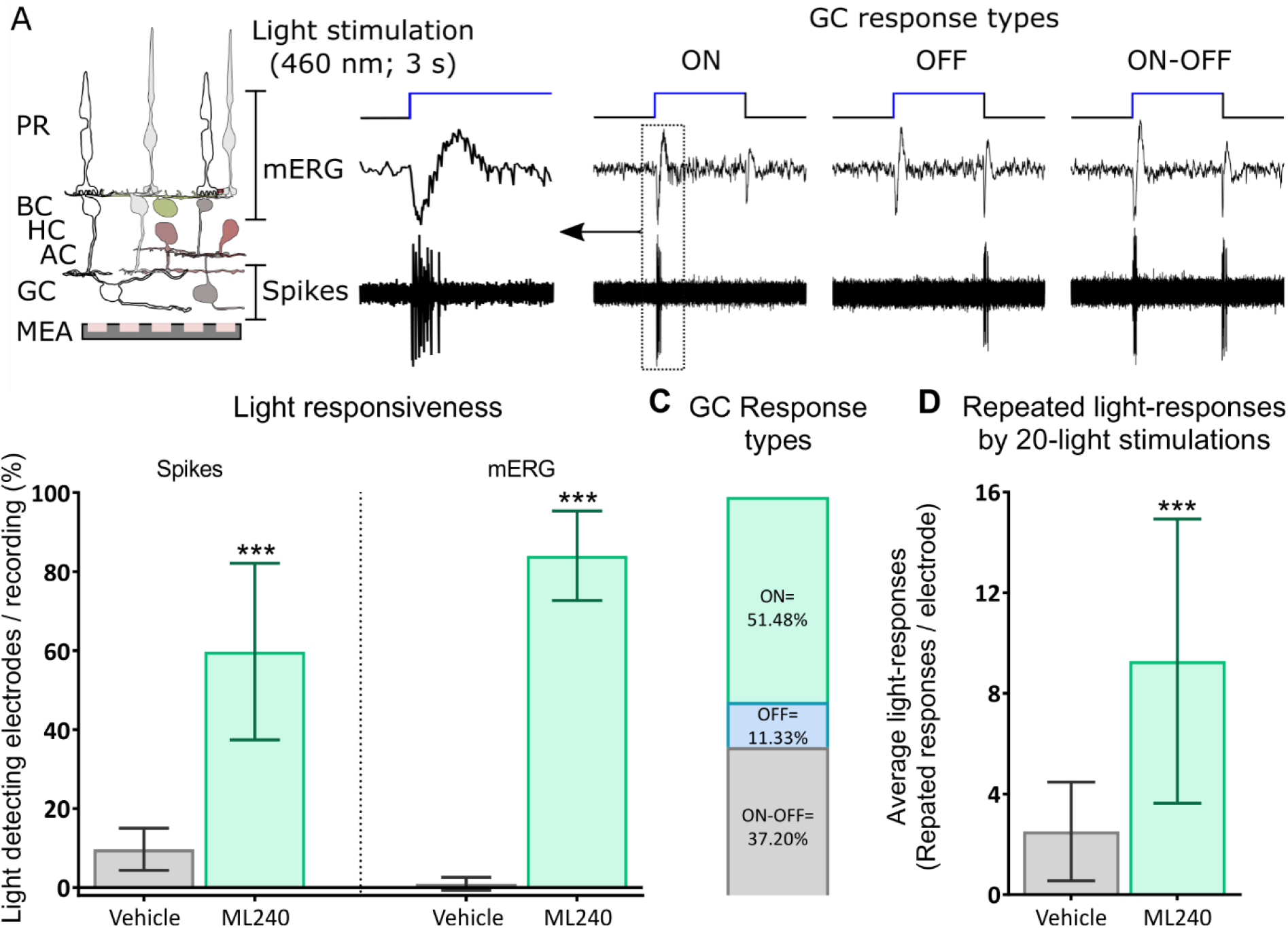
Increased light response of *in vitro* ML240-treated retina. **(A)** Retinal explants (postnatal day 20 and cultivated for 2 days (PN20 DIV2)) treated with ML240 or vehicle were excited using light flashes to distinguish the three ganglion cell types ON, OFF, and ON-OFF, by the light-triggered spike response patterns. On a multi-electrode array (MEA), retinal activity was assessed, recording the inner retinal cell activity (mERG) as well as ganglion cell activity (spikes). Light stimulation: 460 nm wavelength, 35 cd/m^2^ intensity, 3 seconds of light exposure time, and 15 seconds flash interval covering a recording area of 340 μm. **(B)** Light responsiveness of the ML240-treated and vehicle-treated retinal explants recording the number of spikes and mERG signals. The ordinate presents the percentage of electrodes per recording, detecting light-induced activity. **(C)** Percentage distribution of the activated ganglion cell types. **(D)** Light responsiveness of the retinal explants to repeated 20 light flashes (average light response per recording electrode). Data plotted as mean ± SD. Man-Whitney-U test for significance: ***p<0.0001. PR: photoreceptor; BC: bipolar cell; HC: horizontal cell; AC: amacrine cell; and GC: ganglion cell.

To assess the effect of VCP inhibition on retinal function *in vivo*, P23H rats received a single intravitreal injection of ML240 at PN10, and retinal function was assessed by full-field scotopic electroretinogram (ERG) in dark-adapted rats between PN19 and PN21. Light exposure correlates with an increased a-wave corresponding to photoreceptor hyperpolarization, followed by a b-wave response to light corresponding to depolarization of retinal cells post-synaptic to the photoreceptors. In ML240 treated retinae, we observed an increased response to light, as evidenced by higher ERG responses. Both scotopic a-wave and b-wave amplitudes were increased compared to those of vehicle-treated eyes (e.g., 0 log 10 cd s^−1^ m^−2^, a-wave: Vehicle: 46.41 ± 4.63 μV; ML240: 63.11 ± 6.72 μV; p= 0.0102. b-wave: Vehicle: 221.50 ± 20.15 μV; ML240: 281.10 ± 29.40 μV; p= 0.0412 (Fig. 8A)). Putting these data together, the overall effect of ML240 on visual function, determined by scotopic ERGs, indicates an increased light response compared to the vehicle-treated retinae (a-wave: F=10.83, p=0.0013; b-wave: F=7.96; p= 0.0055 (Fig. 8B and C, and Table S2)).

**Figure 8.**
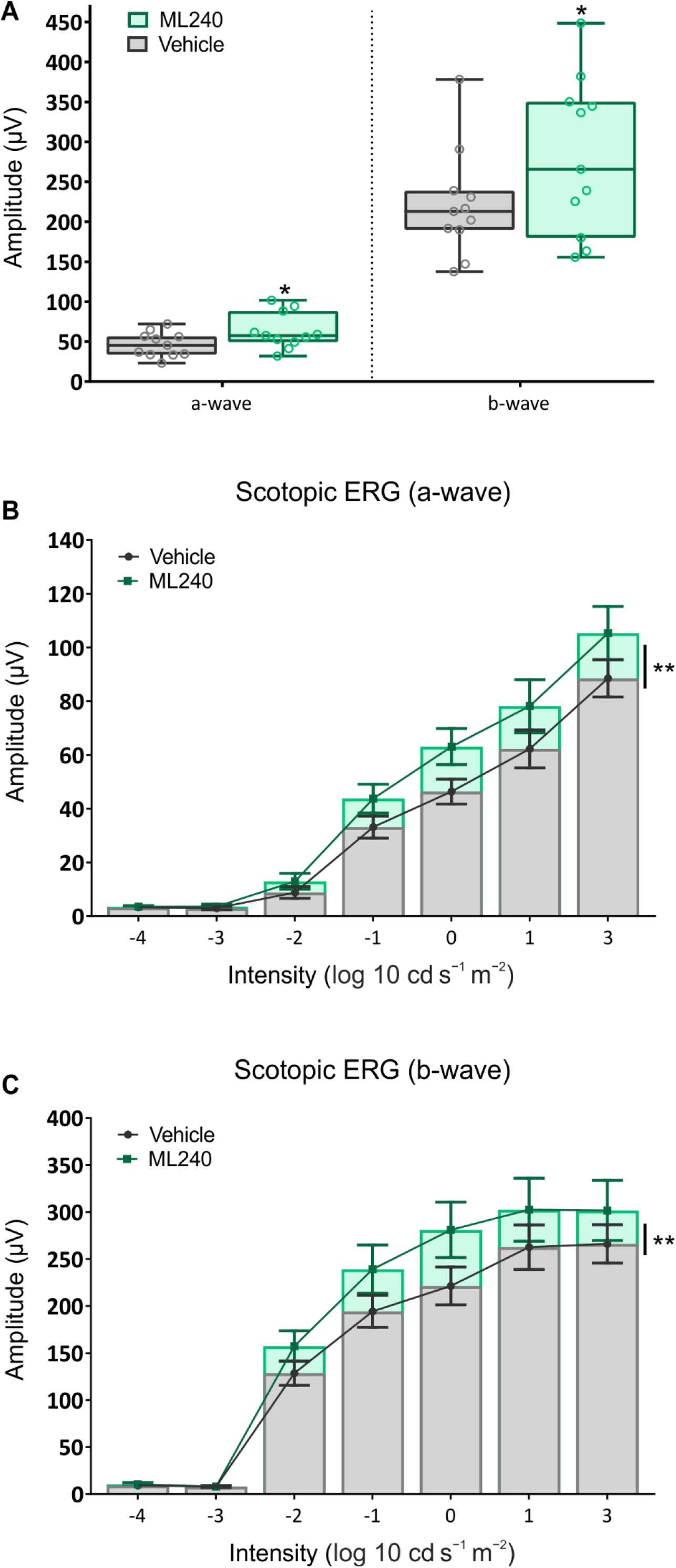
Increased light response of the *in vivo* ML240-treated retina. P23H rats were intravitreally injected at PN10, and scotopic ERG responses performed between PN19 and PN21 (medium-term). **(A)** A-wave amplitude at 0 log cd s^−1^ m^−2^, **(B)** Full-field scotopic a-wave; **(C)** b-wave amplitudes were significantly increased in the ML240 treated eyes. As flash intensities of light were increased, both a- and b-waves increased proportionately in P23H rats injected with ML240. In **(A)** data plotted as means ± minimum to maximum values, n=11, Paired two-sided Student’s t-test. In **(B)** and **(C)** data plotted as means ± SEM, n=11, Two-way ANOVA with Bonferroni multiple comparison test; **p<0.01; *p<0.05.

In conclusion, VCP inhibition by ML240 resulted in improved retinal function correlating with increased cell survival, preserved photoreceptor morphology, and RHO’s corrected distribution.

### The allosteric VCP inhibitor NMS-873 improves the photoreceptor survival and the retinal function in the P23H KI mouse model

Besides competitive VCP inhibitors like ML240, other VCP inhibitors act in an allosteric manner. One of them is the NMS-873, one of the most potent and specific VCP inhibitors described to date, which shows high stability, specificity, and potency (31, 56–59). NMS-873 inhibits both ATPase domains of VCP, whereas ML240 is specific for D2 (19). In order to double-check if the obtained effects are due to VCP inhibition, and following the same experimental rationale, we tested the NMS-873 in a serum-free organotypic culture system, and then we performed *in vivo* intravitreal injections in the heterozygous P23H KI mouse model. P23H KI retinae were isolated from PN14 and cultivated for six days (Fig. 9). Tissues were treated every second day with 5 μM NMS-873, and the vehicle-controls were treated with DMSO.

**Figure 9.**
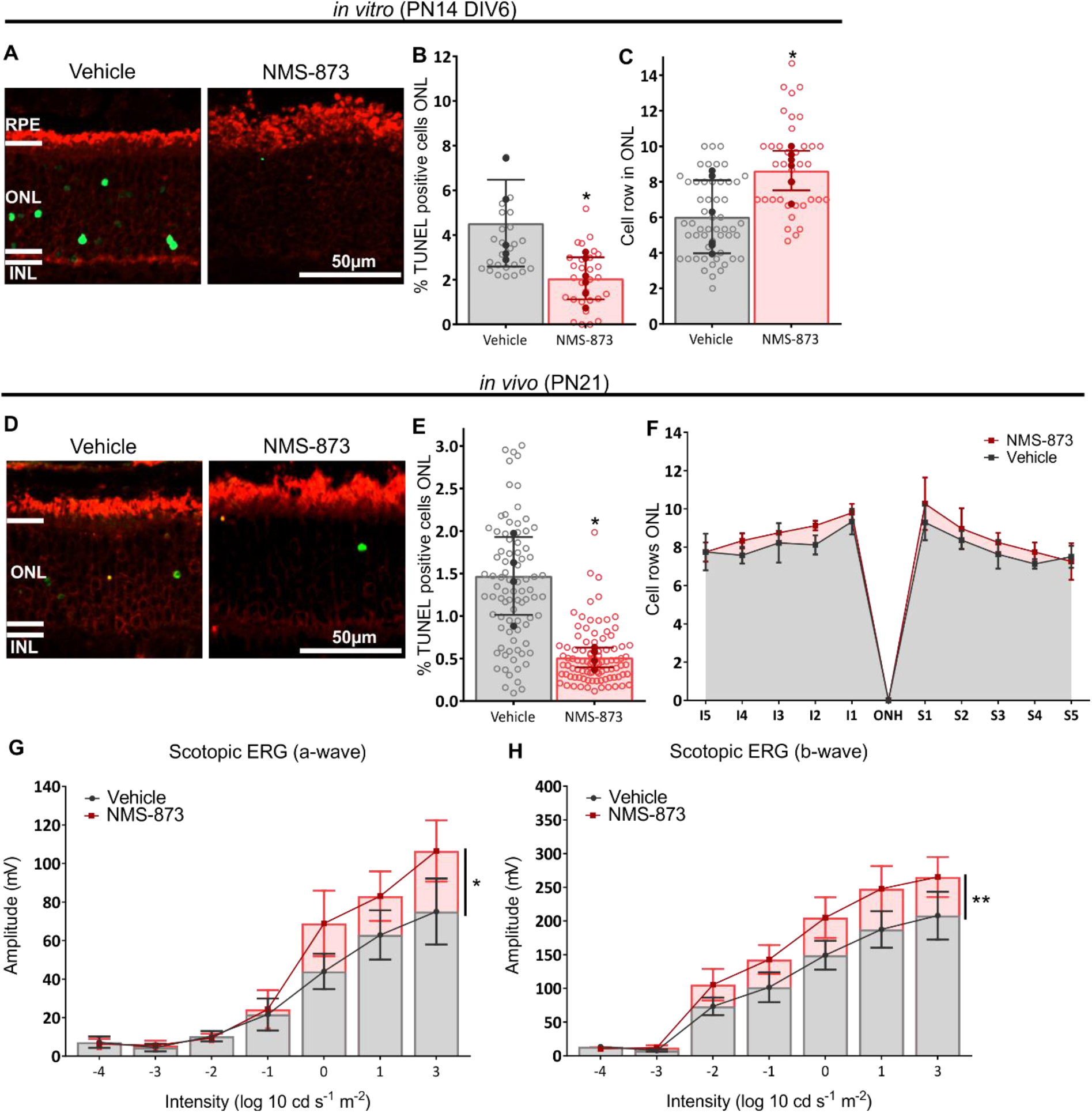
Inhibition of VCP by NMS-873 protects degenerating rod photoreceptors *in vitro* and *in vivo* in P23H KI mice. (**A**) Retinae from P23H KI mice were explanted at postnatal day 14 (PN14) and kept for 6 days *in vitro* (DIV6), and treated every second day with either NMS-873 or DMSO. Explants were stained with TUNEL assay (green) and antibody against RHO (red). NMS-873 treatment restores RHO localization in P23H KI mice retinae explants. Bar chart shows the significantly decreased percentage of TUNEL-positive cells in P23H KI retinae explants treated with NMS-873 compared to the respective DMSO vehicle control. Significant increase in the quantification of the remaining ONL cell rows for the PN20 P23H KI fixed retinae treated with NMS-873 compared to the DMSO vehicle control. Single intravitreal injection in P23H KI mice was performed at postnatal day 11 (PN11) and analyzed at PN21. (**D**) Representative images of the stained retinae using the TUNEL assay to detect cell death (green) and RHO (red) *in vivo.* (**E**) Bar chart shows the decreased percentage of TUNEL-positive cells and (**F**) increased the remaining cell rows *in vivo* (n=4). The full-field scotopic (**G**) a-wave and (**H**) b-wave amplitudes were significantly increased in the NMS-873 treated eyes (n=5). Data plotted as mean ± SEM. (**B**, **C**) Unpaired t-test with Welch correction, n=4. (**E**) paired t-test, n=4. (**F** - **H**) Two-way ANOVA with Bonferroni multiple comparison test; **p<0.01; *p<0.05. RPE: retinal pigment epithelium; ONL: outer nuclear layer; INL, inner nuclear layer. Scale bar: 50μm.

The percentage of TUNEL-positive cells compared to the total number of ONL cell nuclei was significantly reduced by NMS-873 treatment (Vehicle: 4.53 % ± 1.94; NMS-873: 2.06 % ± 0.94 p=0.04 (Fig. 9B)). Thus, the increased survival was reflected in the increased number of remaining cell rows in the ONL (Vehicle: 6.04 rows ± 2.06; NMS-873: 8.63 rows ± 1.12 p=0.026 (Fig. 9C)).

Next, we evaluated the effect of VCP inhibition by NMS-873 in P23H KI mice *in vivo* by intravitreal injection. We selected PN11 as the injection day and PN21 as our evaluation time-point. Similar to our observations *in vitro* and *in vivo* the previous results with ML240 and EerI, we found a significant decrease in the percentage of TUNEL-positive photoreceptor cells after VCP inhibition (Vehicle: 1.47 % ± 0.46; NMS-873: 0.51 % ± 0.12 where p=0.043 (Fig. 9E)), as well as a statistically significant increase in the number of remaining cell rows in the ONL (Fig. 9 F, and Table S3). As seen in ML240 and EerI in the P23H rats, NMS-873 also restored RHO distribution in P23H KI mice *in vitro* and in *vivo*.

Photoreceptor function was assessed by full-field scotopic ERG in dark-adapted P23H KI mice at PN21. Both scotopic a-wave and b-wave amplitudes were increased compared to those of vehicle-treated eyes (a-wave: F=4.22, p=0.0447; b-wave: F=8.69; p= 0.0047 (Fig. 9G and H, respectively, and Table S4).

These results confirm the protective effect of VCP inhibition by three different types of inhibitors and even in two different RP animal models, P23H transgenic rats and P23H KI mice.

## DISCUSSION

VCP has been considered an anti-cancer target, with a VCP inhibitor CB-5083 evaluated in phase 1 clinical trials, NCT02223598 and NCT02243917 (60). Therefore, the pro-survival and neuroprotective effect observed after inhibition of VCP appears counterintuitive at first glance. We report here that VCP inhibition attenuates adRP disease progression in the P23H mediated animal model of RP. In contrast to the effect on cancer cells (i.e., induction of apoptosis), in photoreceptors expressing mutant RHO^P23H^, VCP inhibition significantly increases survival and functionality. VCP inhibition also corrects RHO’s aberrant distribution and restores physiological localization similar to that of the WT phenotype.

Previous studies have revealed that misfolded RHO^P23H^ is retained in the ER and is removed by ERAD (61). VCP interacts with misfolded RHO^P23H^, promoting its retro-translocation and proteasomal clearance (35). In the transgenic *Drosophila Rh1P37H* (the equivalent of mammalian *RHO_P23H_*), inhibition of VCP leads to structural and functional improvements of the insect eye (36). In agreement with these observations, ultrastructural analysis by EM confirmed the improvement of morphology within the OS of treated P23H rats. The typical mislocalization of RHO immunostaining observed in the P23H retinae was almost completely corrected to its physiological localization within OS. However, at this point, it remains unclear whether RHO^P23H^, RHO^WT^, or both are trafficked from the IS to the OS. In *Drosophila*, RHO mutants can recruit WT RHO into intracellular aggregates (62–64). In contrast to that, recent studies suggest that in mammalian cells, misfolded RHO^P23H^ does not aggregate with properly folded RHO^WT^ (65, 66), although other evidence does suggest there is a dominant-negative effect of RHO^P23H^ on the WT protein (2, 67).

We found elongated OS in all groups treated with VCP inhibitors. Since RHO is the most abundant integral membrane protein of OS, representing >90 % of all protein localized in this compartment (5, 6), the observed elongation is likely resulting from enhanced membrane traffic of RHO to the OS and its reduced retention in the ER. VCP inhibition might lead to recovered RHO distribution by several potential mechanisms: either a redistribution of RHO^P23H^, improvement RHO^P23H^ folding or a decrease of RHO^WT^ degradation. Since the two major intracellular protein degradation systems are proteasomal degradation and autophagy, another option is increased degradation of RHO^P23H^ via upregulated autophagy as compensation for the attenuated proteasomal degradation (68, 69). However, further studies are necessary to clarify this.

Previous reports have established a destabilizing effect of misfolded RHO on the disk membranes of OS, potentially mediated by aggregation of the mutated unstable RHO^P23H^. Aggregates formed by RHO^P23H^ could disrupt the correct membrane structure, resulting in vesiculotubular structures and disk breakdown (55, 70). With ML240 treatment, we observed a marked reduction of the vesiculotubular structures in the OS of the P23H rat model retinae. Following Haeri and Knox’s hypothesis, this may result from a change in the RHO^P23H^/RHO^WT^ ratio in the disk membrane. Thus, less RHO^P23H^ and/or increased RHO^WT^ would lead to fewer protein aggregates in the OS and a reduced formation of vesiculotubular structures. Two mechanisms could be involved in altering the balance between mutant and WT protein. As excessive protein retrotranslocation has been described in the *Rh1^P37H^* insect model (36, 68), VCP inhibition may impair this retrotranslocation and consequently increase the amount of RHO^WT^ as well as *Rh1^P37H^* that reaches the OS disk membrane. Alternatively, as VCP is related to selective autophagy modulation (71, 72), rebalancing the autophagic pathway may reduce retinal degeneration caused by protein misfolding (73). In fact, it has been suggested that in the retina of P23H mice, normalization of the autophagic flux relative to proteasome activity may support photoreceptor cell homeostasis, resulting in increased photoreceptor cell survival (74).

The improvements in RHO localization and OS structure mediated by VCP inhibition were accompanied by significant photoreceptor function improvements, as determined by light responsiveness in the retinal explants and ERG responses *in vivo*. The improved photoreceptor function is likely to reflect a combination of enhanced photoreceptor survival and improved photoreceptor homeostasis. At the time points studied, short and medium-term in explants and *in vivo*, the functional improvements observed were greater than the effects on survival, suggesting that photoreceptor homeostasis restoration could be a significant factor contributing to the enhanced light responses. This effect could occur due to a combination of a reduction in an inhibitory effect of misfolded RHO^P23H^ on RHO activation or regeneration and restoration of the OS structure, enabling improved phototransduction. The enhanced photoresponses further illustrate the potential benefit of VCP inhibition as a potential therapy.

Our previous work revealed altered energetic patterns and metabolic failure prior to retinal degeneration in the *Drosophila Rh1^P37H^* model (12). When analyzing the proteome of young *Rh1^P37H^* retinas, we observed a coordinated upregulation of energy-producing pathways and an attenuation of energy-consuming pathways involving the target of rapamycin (TOR) signaling, which was reversed in older retinas at the onset of photoreceptor degeneration. With a combination of pharmacological and genetic approaches, we demonstrated that chronic suppression of TOR signaling (using the inhibitor rapamycin) strongly mitigated photoreceptor degeneration, indicating TOR signaling activation by chronic *Rh1^P37H^* proteotoxic stress is detrimental for photoreceptors. Genetic inactivation of the ER stress-induced JNK/TRAF1 axis and the APAF-1/caspase-9 axis, activated by damaged mitochondria, dramatically suppressed *Rh1^P37H^*-induced photoreceptor degeneration, identifying the mitochondria as mediators of *Rh1^P37H^* toxicity. Thus, the distortion of photoreceptors’ energetic profile leading to a prolonged metabolic imbalance accompanied by the mitochondrial failure may be a driving force for photoreceptor degeneration associated with a *RHO_P23H_* mutation. Therapies normalizing metabolic function could be used to alleviate the imbalance associated with excessive energy wasting in photoreceptor cells. In line with this consideration, it has been shown that inhibition of the ATPase activity of VCP can exert neuroprotection of photoreceptors in an animal model of recessive RP (75, 76), ganglion cell death in a glaucoma model (77), as well as ameliorate retinal ischemia (78). Thus, we hypothesize that the high energy cost necessary for removing misfolded RHO^P23H^ (79) is mitigated by targeting VCP ATPase domains, resulting in stemmed loss of photoreceptors and better retinal functionality.

It is worth noting the sustained protective effect after a single injection of both VCP inhibitors. The usual half-life range for small molecules is between 1–10 hours since small lipophilic compounds are cleared rapidly across the blood-ocular barriers to the systemic bloodstream (80–82). We calculated the expected vitreal half-life for both ML240 and EerI to be approximately 25 minutes, with 98 % being eliminated within 3 hours. Despite this estimated short half-life, both VCP inhibitors remain protective until PN30, 20 days after the single injection. The sustained effect suggests a prolonged mode of action pertaining to photoreceptor cell survival or a mechanism of selective drug retention. In fact, the vitreous and iris can serve as a reservoir for drugs and as a temporary storage depot for metabolites (83, 84). Drug protein binding likely increases retention, given that proteins have a much higher half-life in the vitreous than small molecules (82). EerI acts as an irreversible inhibitor of VCP (38, 85), a fact that could, at least partially, explain a long-lasting effect. However, NMS-873 and ML240 are reversible VCP inhibitors (19, 38), and their sustained protective effect is evident. Thus, it remains unexplained how exactly the drug to target binding or an off-target binding of these compounds contributes to the observed prolonged effect.

We were able to reproduce the neuroprotective achievements using a different kind of inhibitor, NMS-873, and in a second animal model, the P23H KI mice, which more accurately models the human genotypic and phenotypic RHO^P23H^.

Taken together, this study demonstrates a pre-clinical proof of concept that VCP inhibition can effectively attenuate retinal degeneration caused by protein misfolding in mammalian photoreceptors. In humans, retinal degeneration, due to a *RHO_P23H_* mutation, takes on average three decades before legal blindness is reached. This finding opens a decent therapeutic window for a potential clinical application of VCP inhibition, far longer than in the rodent model. Future studies, however, are necessary to assess drug administration’s safety as a primary endpoint, followed by clinical studies to improve visual outcome in retinal degeneration caused by the *RHO_P23H_* mutation in humans.

## METHODS

### Animals

Homozygous P23H transgenic rats (produced by Chrysalis DNX Transgenic Sciences, Princeton, NJ) of the line SD-Tg(P23H)1Lav (P23H-1) were kindly provided by Dr. M.M. LaVail (University of California, San Francisco, CA) or by the Rat Resource and Research Center (RRRC) at the University of Missouri. Animals were housed in the animal facilities of both the Institute for Ophthalmic Research and the University College London, under standard white cyclic lighting, with access to food and water *ad libitum*. To reflect the genetic background of adRP, we employed heterozygous P23H rats obtained by crossing with WT rats (CDH IGS Rat; Charles River, Germany). *Rho^P23H/P23H^* (P23H KI) mice were kindly provided by Dr. K. Palczewski (University of California, Irvine, CA) or purchased from Jackson Laboratory (B6.129S6(Cg)-Rho^tm1.1Kpal^/J, Stock No 017628). The *Rho^P23H/P23H^* mice were crossed with WT C57BL/6J mice to produce P23H heterozygous mice to reflect the genetic background of adRP.

### Organotypic Retinal Explant Cultures of P23H transgenic rats and P23H KI mice

Retinae were isolated with the RPE attached as described previously (41, 86). Briefly, PN9 and PN20 P23H transgenic rats were sacrificed; the eyes were enucleated in an aseptic environment and pretreated with 12 % proteinase K (MP Biomedicals, 0219350490) for 15 minutes at 37 °C in R16 serum-free culture medium (Invitrogen Life Technologies, 07490743A). The enzymatic digestion was stopped by the addition of 20 % FBS (Sigma-Aldrich, F7524). Retina and RPE were dissected, and four radial cuts were made to flatten it. Tissue was transferred to a 0.4 μm polycarbonate membrane (Corning Life Sciences, CLS3412), having the RPE side touching the membrane. The inserts were placed into six-well culture plates and incubated in supplemented R16 nutrient medium at 36.5 °C. The eyes were then randomly assigned to either untreated, vehicle, or VCP inhibitor groups. The VCP inhibitor cultures were treated with either ML240 (20 μM, TOCRIS, Bio-Techne GmbH, 5153) or Eeyarestatin I (10 μM, TOCRIS, Bio-Techne GmbH, 3922). Both compounds were pre-diluted in DMSO (Sigma-Aldrich D2650). DMSO (1% and 0.5%, respectively) was diluted in the culture medium to generate the corresponding vehicle controls. The medium was changed every second day. The PN9 cultures were fixed at DIV6, which correspond to PN15, the peak of degeneration *in vivo* age-matched mutants. The PN20 cultures were fixed at DIV2.

P23H KI heterozygous mice were sacrificed at PN14, and the eyes were enucleated in an aseptic environment. Retinal explants were randomly assigned to either vehicle or 5μM NMS-873 (Xcessbio, M60165-b) treatment. The medium was changed every second day, and retinae cultures were fixed at DIV6, which correspond to PN20, the peak of degeneration *in vivo* age-matched heterozygous P23H KI mice.

### *In vivo* treatment

We divided the transgenic rats into three experimental groups: short-term (ST), retinae analyzed at PN15, medium-term (MT), retinae analyzed at PN21, and long-term (LT), retinae analyzed at PN30. A single intravitreal injection of ML240 or EerI to all P23H rats previously anesthetized (ST and LT: Diethyl ether, Merck Millipore, 100931, or MT: intraperitoneal injection of ketamine (30 mg/kg) and medetomidine (5 mg/kg)) took place at PN10 at the level of the temporal or nasal peripheral retina. Assuming an average rat eye volume of ~25 μl at PN10-30 (45), and an even compound distribution, and in order to achieve the same concentration as in the organotypic cultures (20 μM of ML240 and 10 μM of EerI), the animals received intravitreally 1 μl of 0.4 mM ML240 or 0.5 μl of 0.4 mM EerI on the right eye. The left eye was sham injected with the same volume of vehicle solution and served as contralateral control.

A single intravitreal injection of NMS-873 was performed at PN11 in previously anesthetized (ketamine/medetomidine at 0.08 ml/10 g administered via an IP injection) P23H KI mice in the temporal peripheral retina. Assuming an average mice eye volume, at PN11, is ~5μl (87) and in order to achieve the same concentration as in the organotypic cultures, 5 μM NMS-873, the animals received intravitreally 0.5 μl of 0.05 mM NMS-873 on the one eye while another eye was sham injected with the same volume of vehicle (DMSO) solution. Animals were monitored daily for any adverse effects and for body weight gain. At PN21, mice were subjected to ERG before they were humanely killed by cervical dislocation.

### Assessing the intravitreal clearance of ML240 and EerI

The CL_ivt_ of ML240 and EerI were calculated *in silico* using the QSPR model (43). However, one should notice that ML240 and EerI are found to be more lipophilic than the compounds used in building the model, and thus, the model predictions may not be accurate, but anyway, give an approximation for vitreal drug elimination. The CL_ivt_ value of ML240 and EerI was calculated using the QSPR equation: LogCL_ivt_ = −0.25269–0.53747 (LogHD) + 0.05189 (LogD7.4), where HD is the number of hydrogen bond donor atoms and LogD7.4 is the calculated logarithm of the octanol-water coefficient at pH 7.4 of ML240 and EerI. Since small lipophilic compounds are cleared from the vitreous mainly through the RPE (44), the CL_ivt_ obtained in rabbit eyes was scaled down to rat eyes. Based on the equation CL_ivt_ = P × S, (P: drug permeability in the RPE, S: surface area of the RPE), the CL_ivt_ of small lipophilic compounds in rats was expected to be ~13 times smaller than in rabbits. The RPE surface areas in rats and rabbits are 39 mm^2^ (88) and 520 mm^2^ (89), respectively. Assuming similar permeability of rat and rabbit RPE, the CL_ivt_ in the rat would be 0.04 ml/h (rabbit value: 0.58 ml/h for ML240, and 0.55 ml/h for Eerl). The volume of distribution (Vd) after intravitreal injection is similar to the anatomical volume of the vitreous (43), so that 25 μL for the rats used in the current project (PN10-30) (45). The expected half-life in the vitreous was calculated using the equation: t_1/2_ = ln2 Vd/CL_ivt_.

### Histology

Tissues were immersed in 4 % paraformaldehyde in 0.1 M phosphate buffer (PB; pH 7.4) for 45 minutes at 4 °C, followed by cryoprotection in graded sucrose solutions (10 %, 20 %, 30 %) and embedded in cryomatrix (Tissue-Tek® O.C.T. Compound, Sakura® Finetek, VWR, 4583). Radial sections (14 μm thick) were collected, air-dried, and stored at −20 °C.

#### TUNEL assay

TUNEL assay (90) was performed using an *in situ* cell death detection kit conjugated with fluorescein isothiocyanate (Roche, 11684795910). DAPI (Vectashield Antifade Mounting Medium with DAPI; Vector Laboratories, H-1200) was used as a nuclear counterstain.

#### Immunofluorescence staining and image analysis

Sections were incubated overnight at 4 °C with RHO mAb (Sigma-Aldrich, MAB5316). Fluorescence immunocytochemistry was performed using Alexa Fluor® 568 conjugated secondary antibody (Invitrogen, A-11031). Negative controls were carried out by omitting the primary antibody. The RHO staining’s immunofluorescence intensity was calculated, scoring the mean maximum intensity for a selected central ROI of the ONL of each image within 12 images from three animals per treatment using the Zen 2.3 software. The OS mean length was obtained using the Zen 2.3 software for three selected positions for each image, measuring the OS’s length within 15 (*in vitro*) or 12 images (*in vivo*) from three animals per treatment.

### Transmission Electron Microscopy

Retinal samples were fixed in 2.5% glutaraldehyde, 2 % paraformaldehyde, 0.1 M sodium cacodylate buffer (pH 7.4, Electron Microscopy Sciences, Germany) overnight at 4°C. After rinsing, samples were postfixed in 1 % OsO4 for 1.5 hours at room temperature (Electron Microscopy Sciences, Germany), washed in cacodylate buffer, and dehydrated with 50 % ethanol. Tissues were counterstained with 6 % uranyl acetate dissolved in 70 % ethanol (Serva, Heidelberg, Germany) followed by graded ethanol concentrations up to 100 %, followed by Propylenoxide. The dehydrated samples were incubated in a 2:1, 1:1, and 1:2 mixture of propylene oxide and Epon resin (Serva, Germany) for 1 hour each. Finally, samples were infiltrated with pure Epon for 2 hours. Samples were embedded in fresh resin in block molds and cured 3 days at 60°C. Ultrathin sections (50 nm) were cut on a Reichert Ultracut S (Leica, Germany), collected on copper grids, and counterstained with Reynold’s lead citrate. Sections were analyzed with a Zeiss EM 900 transmission electron microscope (Zeiss, Jena, Germany) equipped with a 2k x 2k CCD camera.

### Ex-vivo light stimulation and activity recordings

P23H rat retinal explants (PN20 DIV2) were taken under dim red light condition (boxed) from the incubator, divided into half across the center, and placed immediately on the recording electrode field of the recording multi-electrode array (MEA) chamber. The recording was performed within the culturing medium, and the recording chamber temperature was set to 37 °C.

#### Retinal activity recording

The MEA system USB-MEA60-Up-BC-System-E (Multi Channel Systems) equipped with HexaMEA 40/10iR-ITO-pr was utilized to record the retinal activity based on 59 recording electrodes. The recordings were performed at 25000 Hz sampling rate collecting unfiltered raw data. The trigger synchronized operation of the light stimulation (LEDD1B T-Cube, Thorlabs) and MEA-recording were controlled by the recording protocol set within the MCRack software (v 4.6.2, MCS) and the digital I/O – box (MCS).

#### Light stimulation

The light stimulation (470 nm LED M470D2, Thorlabs) was applied from beneath the transparent glass MEA guided by optic fiber and optics. A spectrometer USB4000-UV-VIS-ES (Ocean Optics) was employed to determine the intensity of the applied light stimuli (35 cd/m^2^), pre-recordings.

### Electroretinogram

PN19-21 P23H rats or PN21 P23H KI mice were kept overnight in a ventilated light-tight box for dark-adaption. Subsequent procedures were performed under dim red light conditions. Animals were anesthetized, and pupils were dilated using 1.0% tropicamide (Bausch&Lomb). Scotopic ERG was performed using the Celeris-Diagnosys system and Espion software (Diagnosys, LLC, MA). Bilateral electrodes were positioned on rat’s eyes using a liquid gel (Viscotears, Novartis, AG); to produce increasing simultaneous flash stimuli and record retinal activity. For the full-field scotopic ERG, a 7-step protocol was used with the following intensities: 0.0001 (1.03 Hz), 0.001 (1.03 Hz), 0.01 (0.5 Hz), 0.1 (0.1 Hz), 1 (0.05 Hz), 10 (0.04 Hz), 30 (0.04 Hz) cd s^−1^ m^−2^; with short dark-adaption pause in between. For each step, 3 to 10 recordings were displayed and averaged.

### Microscopy and cell counting

All samples were analyzed using a Zeiss Axio Imager Z1 ApoTome microscope, AxioCam MRm camera, and Zeiss Zen 2.3 software in Z-stack at 20x magnification. For quantitative analysis, positive cells in the ONL of at least three sections per group were manually counted. The percentage of positive cells was calculated, dividing the number of positive cells by the total number of ONL cells. Photoreceptor cell rows were assessed by counting the individual nuclei rows in one ONL and averaging the counts.

Graphs were prepared in GraphPad Prism 7.05 for Windows; Adobe Photoshop CS5 and Corel DRAW X5 were used for image processing.

### Statistics

All data, unless otherwise indicated, were analyzed and graphed using GraphPad Prism 7.05 for Windows. P < 0.05 was considered significant.

#### TUNEL counts, photoreceptor row counts, RHO immunofluorescence intensity, and OS length

The evaluation was performed using one-way ANOVA testing, followed by Bonferroni multiple comparisons test. For multiple comparisons in the inferior and superior retinae in the LT group *in vivo*, two-way ANOVA testing, followed by Bonferroni multiple comparisons test, was conducted.

#### MEA-recording

For the electrophysiology data analysis, custom-developed scripts (MATLAB, The MathWorks) were used, if not indicated otherwise. MEA-recording files were filtered employing the Butterworth 2nd order (MC-Rack, Multi-channel systems) to extract the ganglion cell spikes (high pass 200 Hz) and field potentials (bandpass 2 – 40 Hz). The field potentials recorded by the MEA system corresponds to the human ERG as described mERG by (91). The filtered data were converted to *.hdf files by MC DataManager (v1.6.1.0) for further data processing in MATLAB (spike detection and cell-type determination (ON, OFF, and ON-OFF)) as previously described (92, 93). The Man-Whitney-U test was employed to estimate statistical significance.

#### ERG data analysis

For one-to-one comparisons (vehicle-treated vs. treated), a paired Student’s t-test was implemented. For multiple comparisons, two-way ANOVA testing, followed by Bonferonni multiple comparisons test, was conducted for a- and b-wave ERG amplitudes for all stimulus intensities to compare the effect of ML240 or NMS-873 on retinal function.

### Study approval

Procedures were approved by the Tuebingen University committee on animal protection (§4 registrations from 24.04.2013 and AK 15/18 M, and animal permit AK1/13 for P23H transgenic rats, §4 registrations from 12.08.2019 AK 05/19 M for P23H KI mice) and by the UCL Institute of Ophthalmology, London, UK, ethics committee according to the Home Office (UK) regulations, under the Animals (Scientific Procedures) Act of 1986, and performed in compliance with the Association for Research in Vision and Ophthalmology ARVO Statement on animal use in ophthalmic and vision research. All efforts were made to minimize the number of animals used and their suffering.

## Authors contributions

B.A.-G. designed experiments, carried out and treated organotypic cultures, performed intravitreal injections, prepared, stained, and imaged histological samples, analyzed the experimental data, and wrote the manuscript. M.S. carried out and treated organotypic cultures, prepared, stained, and imaged histological samples. M.S. and W.H. planned and carried out the ex-vivo light stimulation and activity recordings of the retinal explants. M.S. and E.M.A. performed the *in silico* calculations. R.G., K.H, K.Z., H.P., R.A., and. M.C. performed intravitreal injections and corresponding ERG and contributed to study planning therein. S.B. performed the EM and prepared histological samples. T.-F.C., R.D., A.U. participated in planning the study, and M.Ue. designed and coordinated the project, participated in designing the experiments, wrote the manuscript, and acquired funding for the studies. All authors have provided feedback on the results, read and approved the final manuscript.

B.A.-G. and M.S. are co-first authors. B.A.-G. is listed first because she contributed more to the conception of the project and the writing of the manuscript.

## Acknowledgments

This study was supported by funds (to M.Ue. and B.A-G) from FFB (Grant PPA-0717-0719-RAD), Kerstan Foundation, European Union’s Horizon 2020 research and innovation programme under the Marie Skłodowska-Curie (Grant agreement No. 722717 – project OCUTHER) and ProRetina Foundation. The personnel of the animal husbandry at the Universitätsklinikums Tübingen and Norman Rieger are acknowledged for the animal care. Christine Henes is acknowledged for her skilled technical assistance with the experiments. Ellen Kilger and Sally Williamson are gratefully acknowledged for language editing and proofreading.

## Notes

The authors have declared that no conflict of interest exists.

### Competing Interest Statement

The authors have declared no competing interest.

