## Supplemental Figure 1 for "Inhibition of VCP preserves retinal structure and function in autosomal dominant retinal degeneration"

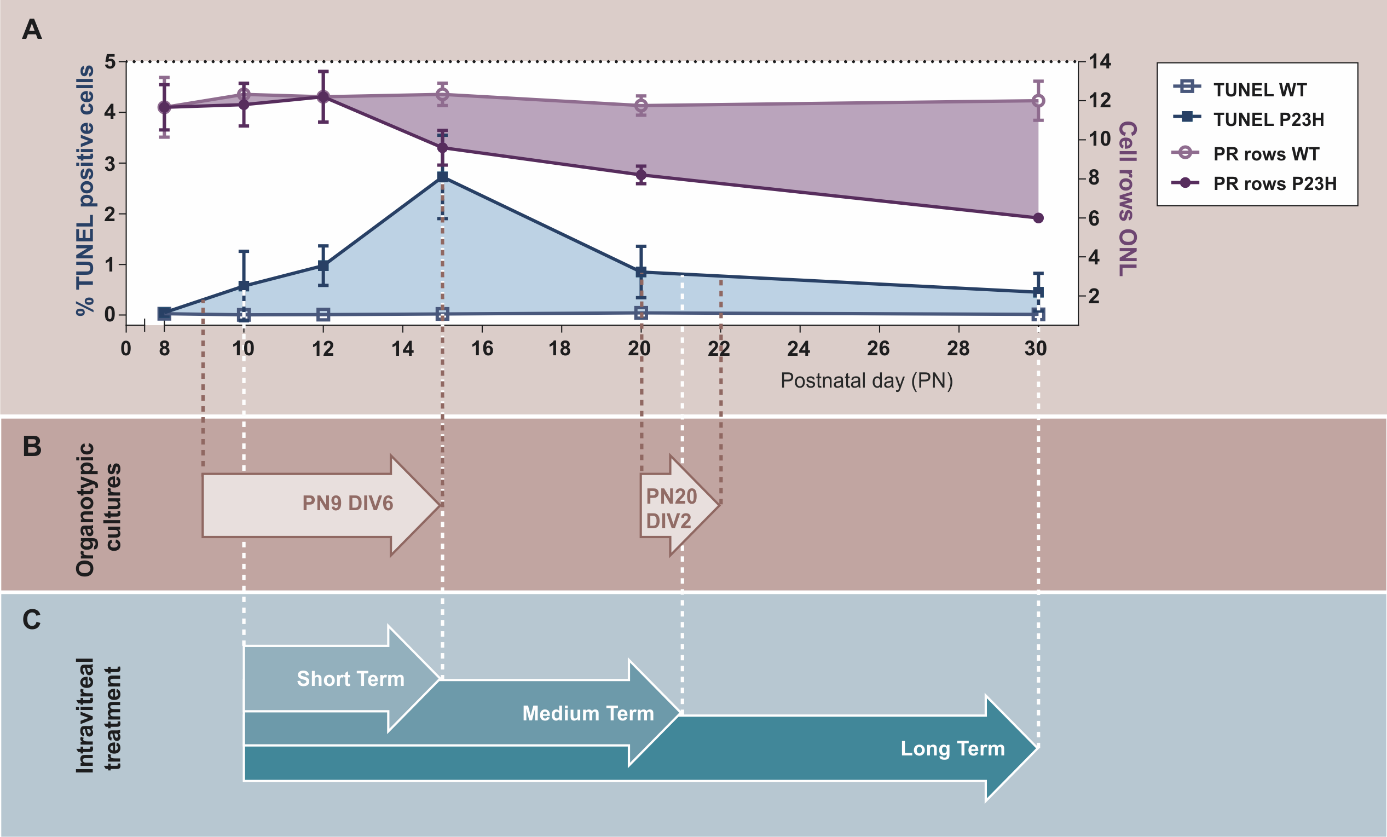

Supplemental Figure 1. **Illustration of photoreceptor cell death progression over the first postnatal month in P23H rats (A) and experimental schedule (B, C)**. **(A)** The diagram displays the increased percentage of TUNEL- positive, dying photoreceptor cells in P23H rats (blue area), showing a peak of cell death at postnatal day 15 (PN15). The loss of photoreceptor cells in P23H rats is reflected in the number of cell rows in the ONL when compared to the WT retina (purple zone); Graphics modified from (Arango-Gonzalez et al., 2014; Kaur et al., 2011). Based on this data, experimental set-ups and design were created: **(B)** Timescale for the organotypic cultures of P23H retinae, explanted at PN9 and cultivated for 6 days (PN9 DIV6) or explanted at P20 DIV2, and **(C)** intravitreal injections of P23H rats at PN10, and then scheduled collection for analyses. The intravitreally injected animals were divided into three groups: short-term (retinae analyzed at PN15), medium-term (retinae analyzed at PN21), and long-term (retinae analyzed at PN30).
