## Supplemental Table S1 for "Inhibition of VCP preserves retinal structure and function in autosomal dominant retinal degeneration"

Supplemental Table S1**. Long Term protective effect of a single intravitreal injection of VCP inhibitors to P23H transgenic rats *in vivo***. Two-way ANOVA analysis with Bonferroni multiple comparison test corresponding to the retinal spidergrams of the inferior and superior hemispheres of treated retinae in Fig. 2 E and F.

|  | ANOVA table | SS | DF | MS | F (DFn, DFd) | P-value |
| --- | --- | --- | --- | --- | --- | --- |
| ML240 | Location | 69.88 | 11 | 6,353 | F (11, 48) = 12.57 | P<0.0001 |
|  | Treatment | 97.51 | 1 | 97,51 | F (1, 48) = 193 | P<0.0001 |
| EerI | Location | 163.30 | 11 | 14,84 | F (11, 144) = 27.99 | P<0.0001 |
|  | Treatment | 93.48 | 1 | 93,48 | F (1, 144) = 176.3 | P<0.0001 |
