## Supplemental Table S2 for "Inhibition of VCP preserves retinal structure and function in autosomal dominant retinal degeneration"

Supplemental Table S2**. Increased scotopic ERG responses of the *in-vivo* ML240-treated retina.** Two-way ANOVA analysis with Bonferroni multiple comparison test corresponding to Fig. 8 B: a-wave amplitudes, and Fig. 8 C: b-wave amplitudes.

|  | SS | DF | MS | F (DFn, DFd) | P-value |
| --- | --- | --- | --- | --- | --- |
| A wave | 3231 | 1 | 3231 | F (1, 136) = 10.83 | P=0.0013 |
| B wave | 33590 | 1 | 33590 | F (1, 136) = 7.963 | P=0.0055 |
