## Supplemental Table S3 for "Inhibition of VCP preserves retinal structure and function in autosomal dominant retinal degeneration"

Supplemental Table S3**. Protective effect of a single intravitreal injection of NMS-873 to P23H KI mice *in vivo***. Two-way ANOVA analysis with Bonferroni multiple comparison test corresponding to the retinal spidergrams of the inferior and superior hemispheres of treated retinae in Fig. 9F.

|  | ANOVA table | SS | DF | MS | F (DFn, DFd) | P-value |
| --- | --- | --- | --- | --- | --- | --- |
| NMS-873 | Location | 49.47 | 9 | 5.497 | F (9, 06) = 10.69 | P<0.0001 |
|  | Treatment | 5.66 | 1 | 5.662 | F (1, 60) = 11.02 | P<0.0015 |
