## Supplemental Table S4 for "Inhibition of VCP preserves retinal structure and function in autosomal dominant retinal degeneration"

Supplemental Table S4**. Increased scotopic ERG responses of the *in-vivo* NMS-873-treated retina.** Two-way ANOVA analysis with Bonferroni multiple comparison test corresponding to Fig. 9 G: a-wave amplitudes, and Fig. 9 H: b-wave amplitudes.

|  | SS | DF | MS | F (DFn, DFd) | P-value |
| --- | --- | --- | --- | --- | --- |
| A wave | 2191 | 1 | 2191 | F (1, 56) = 4.219 | P=0.0447 |
| B wave | 21931 | 1 | 21931 | F (1, 56) = 8.689 | P=0.0047 |
